# Developing simple DNA extraction and PCR-RFLP for *MALE STERILITY 4* (*MS4*) gene in *Cryptomeria japonica* D. Don: Toward an environmentally friendly protocol

**DOI:** 10.1101/2024.08.31.610595

**Authors:** Saneyoshi Ueno, Yukiko Ito, Yoichi Hasegawa, Yoshinari Moriguchi

## Abstract

This study presents the development of a simple DNA extraction method and a novel PCR-RFLP protocol for genotyping the *MALE STERILITY 4* (*MS4*) gene in *Cryptomeria japonica* as a proof of concept. Traditional CTAB-based DNA extraction methods, while effective, involve hazardous chemicals and require high-speed centrifugation, which are impractical in many field settings. Our approach utilizes a household dish detergent-based buffer, sodium chloride, and polyvinylpyrrolidone K-30 to extract DNA from *C. japonica* needle leaf tissues—without the need for liquid nitrogen. The simplicity of this method makes it more accessible and environmentally friendly. The extracted DNA was successfully used in PCR-RFLP analysis, targeting a single nucleotide polymorphism in the *MS4* gene, demonstrating its efficacy for genotyping. The PCR-RFLP markers reliably discriminated between individual genotypes, confirming the practical application of our simple extraction method, even for conifers containing inhibitory substances. This technique is particularly advantageous for use in arboretums and field stations, where the use of hazardous chemicals, specialized equipment, and a draft chamber is limited. Our study contributes to genetic resource management by providing an easy, reliable, and safer method for DNA extraction and genetic analysis.

## Introduction

Nucleic acid extraction is a starting process for genetic analysis and has been a topic of research for many years (Yadav et al. 2018; Dairawan and Shetty 2020; Schenk et al. 2023). The DNA extraction from plant tissue is difficult compared to that from animal tissue, because it contains poly-saccharide, polyphenolics and other secondary metabolites that prevent nucleic acid extraction. This challenge is particularly pronounced in trees, especially conifers like *Cryptomeria japonica*, which are known for their high levels of polyphenolics and polysaccharides. These compounds interfere with DNA extraction and PCR amplification, making the process more complex than with other plant species.

Developing a method that can effectively extract DNA from such challenging plant material is crucial for advancing genetic studies in these species. In 1980, Murray and Thompson (Murray and Thompson 1980) published a DNA extraction method using CTAB (cetyl-trimethylammonium bromide) which efficiently removes polysaccharides from plant tissue. Currently the CTAB-based method is used to extract DNA/RNA from many plant species with wide downstream uses for restriction digestion and PCR experiments without further purification (Doyle and Doyle 1987; Le Provost et al. 2007). One drawback of the method is the use of hazardous chemicals such as chloroform and 2-mercaptoethanol, which are recognized internationally as hazardous substances. Specific handling must comply with the regulations and guidelines of each country. In addition to the use of hazardous chemicals, the CTAB protocol uses centrifugal forces of up to 20,000 g. Unfortunately, arboretums and field stations are rarely equipped with such centrifuges.

One of the simplest methods to extract DNA is the use of chelating resin, InstaGene (Bio-Rad Laboratories, Inc.), which requires only heating the tissue with the resin matrix for DNA extraction.

Tsuruta et al. (2021) used InstaGene Matrix to extract DNA from somatic embryogenic cells of *C. japonica* for marker assisted selection of *MS1* (*MALE STERILITY 1*) gene (Tsuruta et al. 2021). The method was further applied for leaf, pollen, and inner bark tissue of *C. japonica* from which DNA was extracted for *MS1* genotyping (Ueno et al. 2022). Although the InstaGene extraction is simple, the amount of tissue (5–10 mg leaf) used with InstaGene Matrix is a key factor for the success of DNA extraction (Ueno et al. 2022). Too much tissue results in a viscous solution, leading to failure in PCR. Moreover, the process of trial and error with InstaGene (e.g., optimizing the amount of InstaGene matrix used, the duration of heating the samples with the InstaGene matrix, and the PCR thermal profile) can be a time-consuming task, especially for those inexperienced in DNA experiments.

In this study, we aim to develop a simple DNA extraction method and PCR-RFLP (polymerase chain reaction-restriction fragment length polymorphism) genotyping technique for the recently identified *MS4* (*MALE STERILITY 4*) gene in *C. japonica* (Kakui et al. 2023). The mutation in *MS4* involves a single nucleotide polymorphism (SNP) that results in an amino acid substitution. Although a KASP (Kompetitive Allele Specific PCR) (Dipta et al. 2024) marker for the *MS4* gene was recently developed (Watanabe et al. 2024), its application requires specialized equipment such as a real-time PCR machine, which may not be readily available in arboretums or field stations. Another potential approach is developing allele-specific PCR markers, but this method requires extensive optimization of PCR conditions, even though Locked Nucleic Acids (LNA) are included in PCR primers (Latorra et al. 2003), making it more challenging and time-consuming. Therefore, we focused on developing a PCR-RFLP technique that serves as a practical example of applying our simple DNA extraction method. This approach, which avoids the use of hazardous chemicals, aims to provide an accessible, environmentally friendly tool for genetic analysis, offering a versatile solution for researchers in various settings.

## Materials and Methods

Needle leaves from five *C. japonica* individuals (Table 1) were sampled at Niigata Prefectural Forest Research Institute on July 22, 2022, and kept refrigerated at 4 °C until DNA extraction. One needle leaf (approximately 5 mg) was ground in 1 mL of dish detergent buffer (10 % dish detergent ‘Yashinomi Hi-power’ (Saraya Co., Ltd.), 10 % NaCl, and 5% polyvinylpyrrolidone K-30 (PVP) (NACALAI TESQUE) at room temperature using mortar and pestle (Fig. 1). The dish detergent contained 32 % surfactant (sodium alkyl ether sulfates, alkyl betaines, and fatty acid alkanolamides), auxiliary cleaning agents, and stabilizers. The resulting ‘green juice’ (approximately 500 µL) was transferred to a 1.5 mL tube, centrifuged for three minutes with a CHIBITAN-R microcentrifuge (Hitachi Koki, Co. Ltd.) at the maximum speed of 6,200 rpm, and 300 µL of supernatant was then collected in a new 1.5 mL tube. Twice the volume (600 µL) of cold 100 % ethanol was added to the tube and mixed with inverted shaking. The solution was centrifuged for 10 min at room temperature using the CHIBITAN-R, and the supernatant was discarded by decantation. 600 µL of 70 % ethanol was then added to the tube to wash the precipitate, spun briefly by CHIBITAN-R, and discarded by decantation. The precipitate was air-dried at room temperature, and dissolved in 100 µL of TE buffer (NACALAI TESQUE). The DNA concentration was measured by Qubit Fluorometer (Thermo Fisher Scientific).

**Table 1.** Japanese cedar samples used in the current study.

| No<sup>#</sup> | ID | Male flower phenotype | *MS4* genotype<sup>§</sup> |
| --- | --- | --- | --- |
| 1 | Shindai 8 | Sterile | *ms4/ms4* |
| 2 | S8HK5 | Fertile | *Ms4/ms4* |
| 3 | Kamikiri Niigata-58 | Fertile | *Ms4/ms4* |
| 4 | Higashikanbara-5 | Fertile | *Ms4/Ms4* |
| 5 | S8 × Gosenshi-1 | Fertile | *Ms4/ms4* |
| 6 | Setsugai Niigata-1 | Fertile | *Ms4/Ms4* |
| 7 | Kitakanbara-1 | Fertile | *Ms4/Ms4* |
| 8 | Y938 | Fertile | *Ms4/Ms4* |

<sup>#</sup> For development of the PCR-RFLP protocol, trees No 1–4 were used, while for development of the simple DNA extraction method, trees No 1 and 4–7 were used. Tree No 8 was used to test the reproducibility of the protocol.
<sup>§</sup> Genotypes were known previously for trees No 1–5 (Miyajima *et al.* 2010; Hasegawa *et al.* 2018; Hirayama *et al.* 2021), while they were unknown for trees No 6–8 before the present study.

**Fig. 1.**
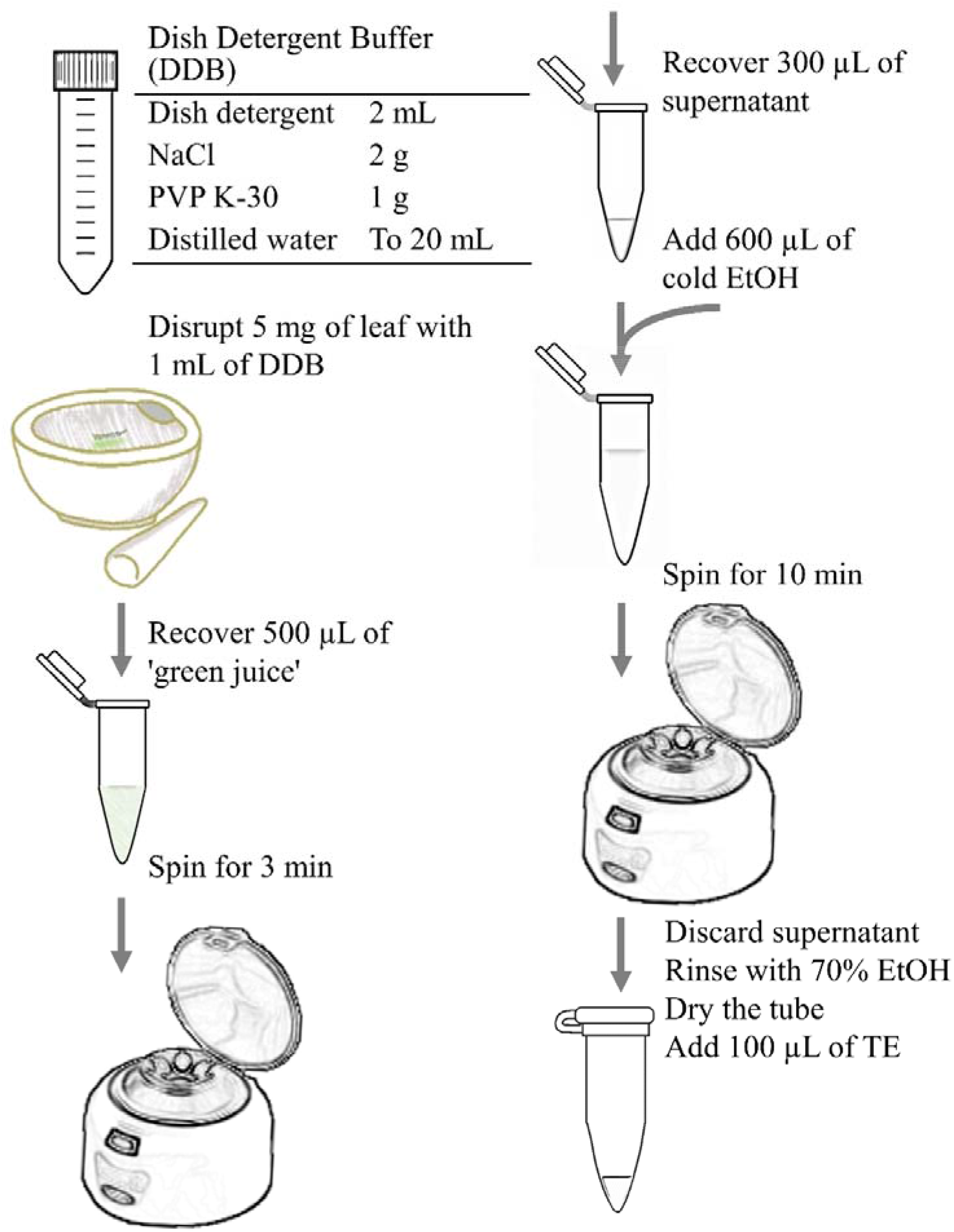
Workflow of the simple DNA extraction method.

In order to demonstrate the practical application for the extracted DNA, a PCR-RFLP protocol was developed to genotype the male-sterile causative mutation of *MS4* in *C. japonica* (Fig. 2) as a model system. PCR primer pairs to amplify the target region were designed by Primer3Plus (Untergasser et al. 2012). The SEQUENCE_TEMPLATE contained the CJt034360 sequence (DDBJ accession number: ICQT01013259) with the *PRIMER_MIN_SIZE* 23, *PRIMER_OPT_SIZE* 23, *PRIMER_MAX_POLY_X* 2, *PRIMER_PAIR_MAX_DIFF_TM* 2, and SEQUENCE_TARGET 301,1 (target causative SNP coordinate (301) along the *SEQUENCE_TEMPLATE*, and the length of 1). The enzymatic function of *MS4* is TKPR1 (tetraketide alpha-pyrone reductase 1), the dysfunction of which causes male sterility in not only *C. japonica*, but also *Arabidopsis* (Grienenberger et al. 2010) and rice (Xu et al. 2019). Restriction enzymes to discriminate the mutant and wild type alleles in *MS4* were searched using the REHUNT program (Cheng et al. 2018). Four samples (Table 1), whose *MS4* genotypes were established elsewhere by artificial crossing (Miyajima et al. 2010; Hasegawa et al. 2018; Hirayama et al. 2021) and whose DNA was extracted using a modified CTAB method (Hasegawa et al. 2020), were used in the development of PCR-RFLP protocol. The PCRs were carried out in a 10 µL reaction containing 1 µL (about 10 ng) of template DNA, 1 µL of each of primer (2 µM stock solution; forward: AGCCATGAATGGAACAACTGTGT, and reverse: GGAGGAAGGCTCGGTCCTATTAT), resulting in a final concentration of 0.2 µM for each primer, and 5 µL of QIAGEN Multiplex PCR Master Mix (Qiagen). The thermal profiles used were as follows: the initial denaturation at 95 °C for 15 minutes, followed by 40 cycles of 95 °C for 30 sec, 60 °C for 90 sec, and 72 °C for 60 sec with the final extension of 72 °C for 10 min. The resulting products (5 µL) were digested with the restriction enzyme *Hae*III (NIPPON GENE) in a 10 µL of reaction containing 2–5 µL of PCR products, 1 µL of 10× Buffer M (resulting in a final concentration of 1×), and 0.1 µL of *Hae*III (high concentration: 30 – 100 units/µL) and incubated for at least three hours or overnight at 37 °C. The PCR products and restriction-digested products were run on 2 % agarose gels, stained with ethidium bromide, and visualized on a UV transilluminator.

**Fig. 2.**
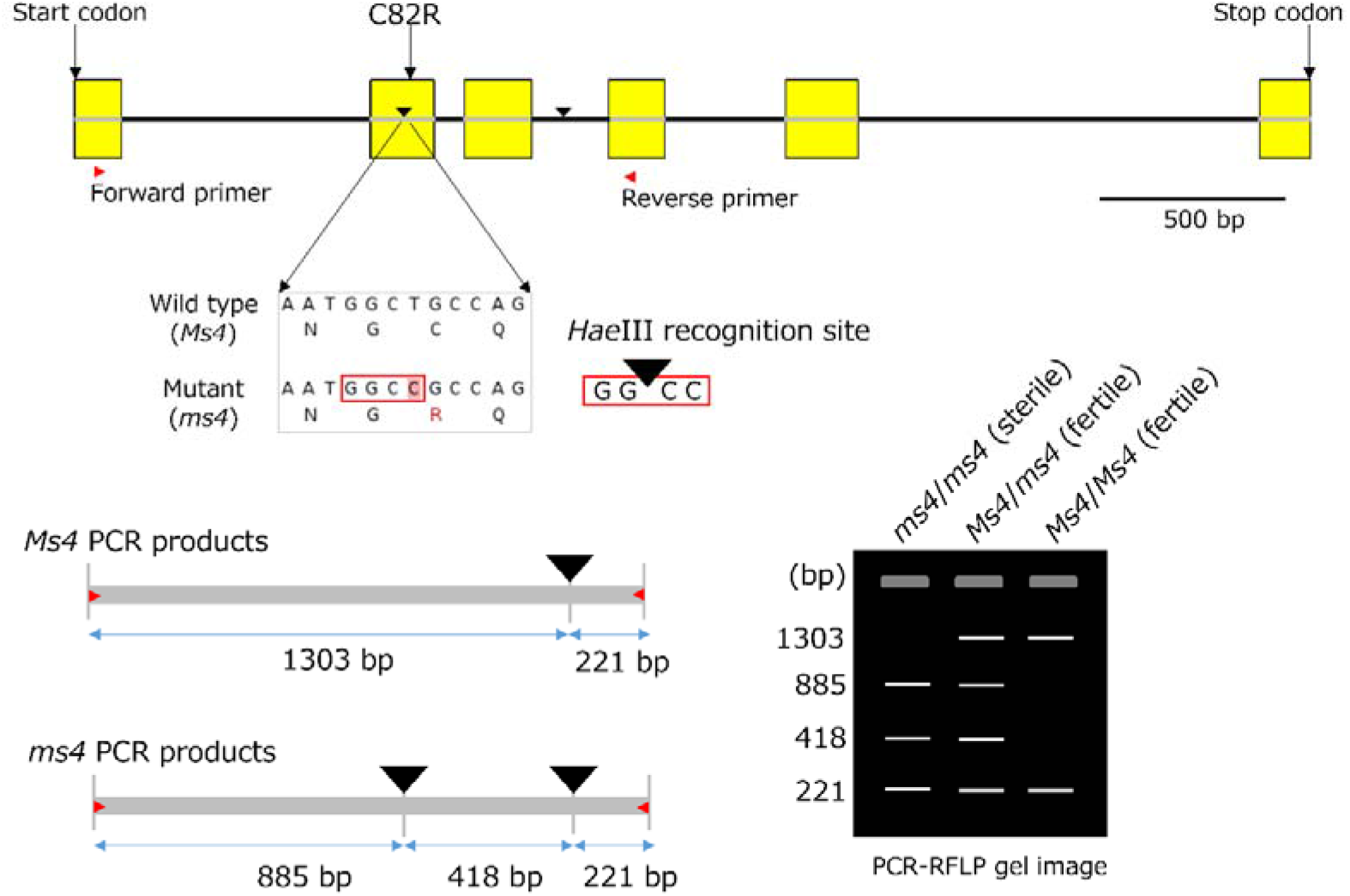
Schematic representation of *MS4* (*TKPR1*) and PCR-RFLP in *Cryptomeria japonica*. Yellow rectangles indicate CDS (coding sequences), which are divided by intron sequences shown by black lines. *MS4* consists of five exons and introns with a total of 3,323 bp from the start to the stop codon. The CDS is translated into 333 amino acid residues, the 82^nd^ of which is cysteine (C) and arginine (R) for the wild type and the mutant, respectively. This is the causative mutation of male sterility by *MS4*. The restriction enzyme, *Hae*III, can recognize the GGCC, the mutant nucleic acid sequence, while the wild type sequence cannot be cut by the enzyme. The PCR-RFLP primer pairs are shown by red triangles; the PCR products from *ms4* allele contain two *Hae*III recognition sequences shown by black triangles, the C82R site, and that in the third intron common with the products from the wild type allele. The resulting electrophoretic profile, with gel wells in gray, shows thee, four, and two bans for *ms4*/*ms4, Ms4*/*ms4*, and *Ms4*/*Ms4* genotype, respectively

Reproducibility of this simple DNA extraction and PCR-RFLP protocol was confirmed with current year needles sampled from a single individual, ‘Y938’ (Table 1) growing at Forestry and Forest Products Research Institute, Tsukuba, Japan on October 11, 2024.

## Results and Discussion

The DNA extracted by the simple method yielded concentrations ranging from 1.01 to 2.11 ng/µL with an average of 1.66 ng/µL (N=8). The number of needles required to reach a sample weight of 5 mg varied, ranging from one to three needle leaves, depending on the individual sample. The PCR-RFLP protocol developed for *MS4* successfully discriminated the individual genotypes (Fig. 3). When DNA extracted by the dish detergent buffer was used for this protocol, expected results were also obtained, demonstrating the successful application of the simple DNA extraction method in this study. We confirmed the reproducibility of this simple DNA extraction and PCR-RFLP protocol by conducting eight independent DNA extraction and PCR-RFLP experiment using the same single individual (Fig. 3). In addition, one of the co-author (Ms. Ito), a beginner of DNA work, has already genotyped over 100 samples with a failure rate of less than 5%. This suggests that the method is not only reproducible but also robust across different operators. If the sample is sufficiently ground, a white precipitate is visible at the bottom of the tube after centrifugation during ethanol precipitation, indicating successful DNA extraction. Even if the precipitate is not visible, the target fragment (1.5 kb) was still typically amplified by PCR. These results indicate that the method is reproducible and robust across a wide variety of samples and operators. Furthermore, using a homogenizer can improve stability and increase DNA yield. Following the same protocol, the Multi-beads Shocker (MB3000; Yasui Kikai Corp., Osaka, Japan) at 2500 rpm for 20 sec achieved a concentration of 5 ng/µL, with a total yield of 500 ng. This demonstrates the method’s potential for high-throughput applications, making it suitable for handling large numbers of samples efficiently without liquid nitrogen.

**Fig. 3.**
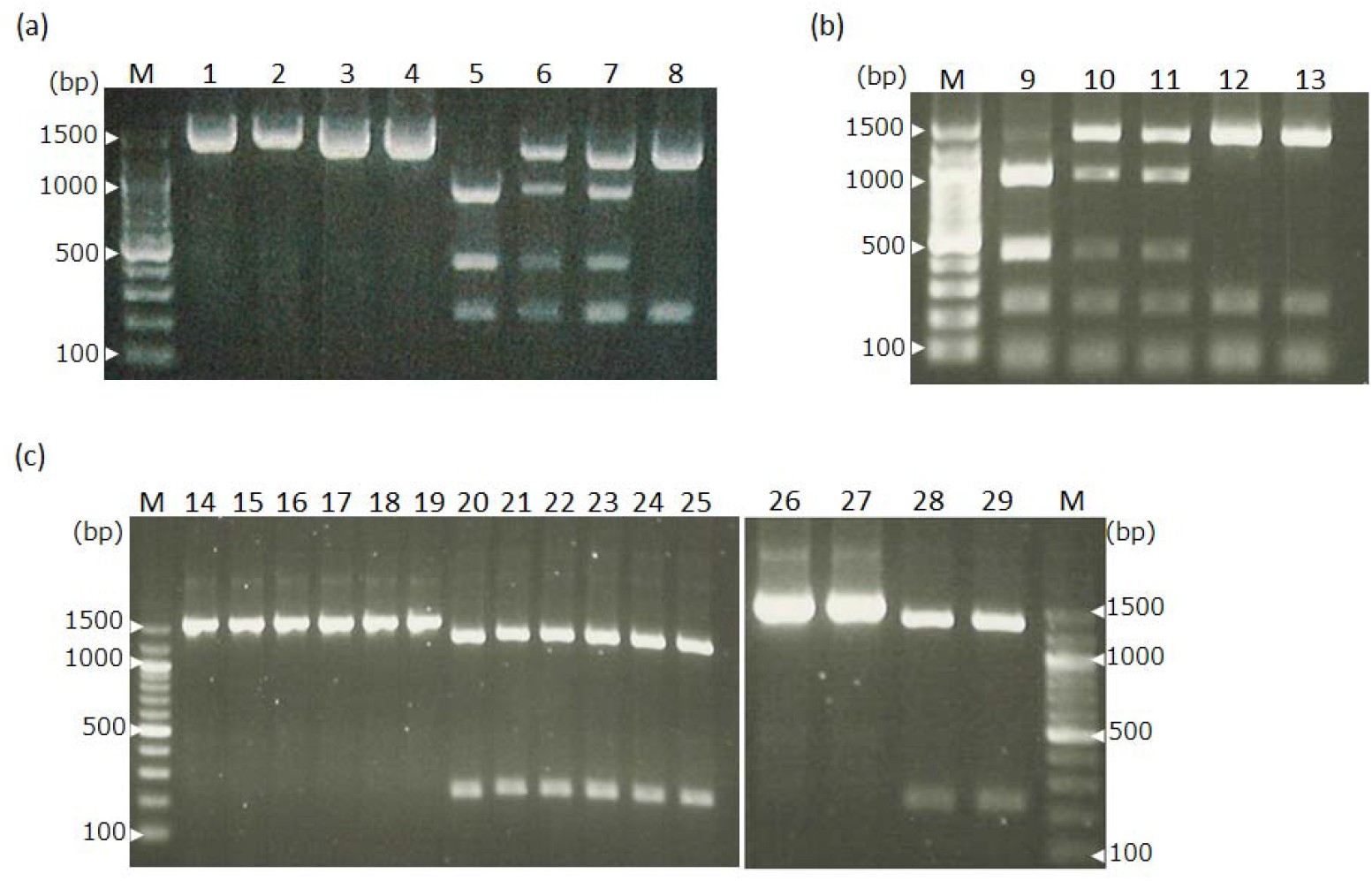
Photographic images for agarose gel electrophoresis. (a)Development of PCR-RFLP for *MS4* gene: PCR products (lanes 1–4) and *HaeIII*-digested products (lanes 5–8) for ‘Shindai 8,’ ‘S8HK5,’ ‘Kamikiri Niigata-58,’ ‘Higashikanbara-5’ with genotypes of *ms4*/*ms4, Ms4*/*ms4, Ms4*/*ms4*, and *Ms4*/*Ms4*, respectively. (b) DNA extracted by the simple extraction using the dish detergent buffer was used for PCR-RFLP genotyping. *Hae*III-digested products for ‘Shindai 8,’ ‘Kamikiri Niigata-58,’ ‘S8 × Gosenshi-1,’ ‘Setsugai Niigata-1,’ and ‘Kitakanbara-1’ are in lanes 9–13. The smallest bands (less than 100 bp) may result from primer dimers or unspecific amplification products. (c) Reproducibility was tested by independently extracting DNA eight times from the same sample individual (‘Y938’) using the dish detergent buffer, followed by PCR-RFLP analysis: PCR products (lanes 14–19 and 26–27) and *HaeIII*-digested products (lanes 20–25 and 28–29). In all (a), (a)and (c), 100 bp ladder was run on the lane M

Because DNA extraction is a basic starting experiment of molecular biology, biology classes in secondary education conduct DNA extraction experiments using commercially available detergent with simple procedures (Ogata et al. 2010; de Andrade et al. 2024). Although these classes commonly use soft tissue plant materials including broccoli flower buds and strawberries, no reports were found for DNA extraction from coniferous plants using dishwashing detergent. In the current study, we developed a simple DNA extraction method with commercially available dishwashing detergent for use in marker assisted management of arboretum or field stations, where the use of hazardous chemicals is not preferable and powerful centrifuges are not available. When DNA extracted with this simple detergent buffer was run on agarose gels, DNA bands were rarely observed. This is probably due to the small amount of DNA extracted that is often fragmented. However, PCR successfully amplified the 1.5 kb fragment, though the amplification of primer dimers or non-specific amplification occurred (Fig. 3).

While polymerases, such as MightyAmp DNA polymerase (Lu et al. 2012), may be suitable for use with crude DNA that contains PCR inhibitors, this study used HotStartTaq DNA polymerase in Multiplex Kit (Qiagen) which is widely used for genetic analysis with microsatellite markers for clonal identification and parentage analysis. Additionally, since the DNA extracted using this method could also be utilized with restriction enzymes, which are generally more sensitive to inhibitors than polymerases in PCR, it suggests that the quality of the extracted DNA is high enough for various molecular biology applications.

For next-generation sequencing applications, this DNA extraction method shows potential compatibility with amplicon sequencing, such as MIG-seq (Suyama and Matsuki 2015). Although the yield and quality may be limiting factors for RADseq (Baird et al. 2008; Peterson et al. 2012), exploring its application to RADseq could be of interest and presents a challenge for future research.

The DNA extraction buffer used in this study (Fig. 1) comprised commercially available dishwashing detergent, NaCl, and PVP, each contributing to the effectiveness of the extraction process. The dishwashing detergent acts as a surfactant, breaking down cell membranes and denaturing proteins, thus facilitating the release of DNA into the solution. There are several brands of dishwashing detergent in the market. Although each product has different ingredients, comparative studies have indicated that any dishwashing detergent could extract DNA (Ogata et al. 2010). Because the concentration of detergent is different for each brand, preliminary experiments to determine the most effective concentration of detergent is necessary. In the current study, we initially tried to use 1 µL of ‘green juice’ extracted by an extraction buffer containing only 2 % dish washing detergent, directly as a template DNA for PCR. Although 1.5 kbp PCR products were successfully obtained, we failed to digest the PCR products by *Hae*III restriction enzyme, probably due to inhibiting contaminants in the ‘green juice.’ We therefore included salting-out/salting-in for proteins, and ethanol precipitation for DNA in the protocol (Fig. 1).

NaCl is indispensable to precipitate DNA by ethanol, while it is expected to work as salt to prevent proteins from dissolving in or precipitating from the buffer. Instead of using dishwashing detergent, laundry detergent may be used (Nasiri et al. 2005). However, measuring a specific amount of detergent powder and dissolving it into a buffer may be more tedious than using liquid type detergent, though any commercially available products commonly found in the laboratory could be a key factor in terms of cost and convenience. The PVP, the other ingredient in the extraction buffer (Fig. 1), was included, because it efficiently removes phenolic compounds, which interact with DNA and prevent DNA extraction (Heikrujam et al. 2020). Extraction buffer without PVP led to failure in *Hae*III restriction digestion in the current protocol for *C. japonica*, though PCR was successful (data not shown). If samples contain less phenolic compounds or only PCR application is needed, PVP could be removed from the extraction buffer.

In conclusion, we successfully developed a simple and effective DNA extraction method that is applicable to conifers, a plant group traditionally difficult to process due to inhibitory substances. The successful application of this method in genotyping the *MS4* gene in *C. japonica* suggests its potential utility in a wide range of plant species, making it a valuable tool for genetic analysis in settings where access to specialized equipment is limited. This approach broadens the scope of molecular biology applications and offers a practical solution for DNA extraction in diverse and challenging environments.

## Acknowledgments

This research was supported by Bio-oriented Technology Research Advancement Institution (BRAIN) Grant (JPJ007097; Project ID 28013B) and FFPRI Grant (#201421, #201906). Samples used in the simplified DNA extraction method were collected by Mr. Iwai of the Niigata Prefectural Forest Research Institute. The authors would like to thank Enago (www.enago.jp) for the English language review.

## Data Archiving Statement

The sequence data used in this study are archived in the DDBJ under the accession number ICQT01013259.

